# Acellular Cartilage-Bone Allografts Restore Structure, Mechanics, and Surface Properties In Vivo, but Limit Recellularization and Integrative Repair

**DOI:** 10.1101/2022.10.27.514135

**Authors:** Jeanne E. Barthold, Luyao Cai, Kaitlin P. McCreery, Kristine Fischenich, Kevin Eckstein, Virginia Ferguson, Nancy Emery, Gert Breur, Corey P. Neu

## Abstract

The repair of articular cartilage after damage is challenging, and clinical interventions to promote regeneration remain elusive. The most effective treatment for cartilage defects utilizes viable osteochondral allografts from young donors, but unfortunately suffers from severe source limitations and short storage time. Decellularized tissue offers the potential to utilize native tissue structure and composition while also overcoming source limitations, but the long-term efficacy of acellular allografts is unknown. Here, we show that acellular osteochondral allografts improve functional and integrative cartilage repair in defect regions after 6 months in a preclinical (sheep) animal model. Functional measures of intratissue strain and structure assessed by MRI demonstrate similar biomechanical performance between implants and native cartilage. Compared to native tissue, the structure, composition, and tribology of acellular allografts conserve surface roughness and lubrication, native cartilage material properties under compression and relaxation, and compositional ratios of collagen:glycosaminoglycan and collagen:phosphate. However, while high cellularity was observed in the integration zones between native cartilage and acellular allografts, recellularization throughout the chondral implant was largely lacking, potentially limiting long-term cellular maintenance in the graft and repair success. Our results advance a suite of joint-to-cellular functional assays, demonstrate the biomechanical efficacy of acellular allografts for at least six months *in vivo*, and suggest that long-term implant success may suffer from a lack of cell migration into the dense decellularized chondral tissue.

## I. INTRODUCTION

Articular cartilage is an essential tissue within the joint that functions as a load-bearing and low-friction surface to enable locomotion. Injury or damage to articular cartilage does not spontaneously heal and can lead to degeneration and osteoarthritis (OA). The loss of cartilage due to disease or overuse leads to an inability for the remaining tissue to maintain mechanical function properly when exposed to daily activities. In the United States, over 32 million people suffer from osteoarthritis of the knee and the majority of those affected experience a lower quality of life and loss of function due to the associated knee pain^1^. If left untreated, injury sites in the cartilage often progress to advanced joint osteoarthritis ^2^. While there are medical treatments and surgical approaches to ameliorate the pain and loss of function, the most likely option to completely treat the condition and restore function at late-stage OA is a total knee replacement using metal and polymer components. The most robust knee replacements typically only last 15-20 years ^3^, which is problematic since over 50% of patients requiring treatment for osteoarthritis of the knee are under the age of 65 ^4^.

A significant hurdle for patients suffering from osteoarthritis is the lack of effective therapies to address cartilage defects leading to osteoarthritis and minimize the need for more invasive replacement surgeries. Two of the most successful current clinical techniques for tissue repair are osteochondral (bone and cartilage) autologous transplantation and osteochondral allograft transplantation. Autologous transplantation involves taking cartilage tissue from a non-load bearing region of a patient’s knee and moving it to the defect region. While long term follow-up studies show success ^5^, this procedure is limited by the defect size that can be repaired, as tissue is sourced from a different part of the knee and can lead to donor site morbidity and associated complications. Osteochondral allografts, or transplant of osteochondral tissue from healthy cartilage of a recently deceased donor, is the most successful therapy to date to delay osteoarthritis progression, and thus a full knee replacement, showing an 82% survival rate after 10 years ^6^. Unfortunately, the quantity of osteochondral allografts available is limited by reliance on healthy donor cartilage and the need for immediate harvest and implant into recipient for optimal success ^7^.

Considering the increase in patients with OA ^8^, there exists a substantial need for cartilage defect therapies that (1) delay OA progression and a total joint replacement, (2) can be delivered in one surgery, (3) have a long shelf life, and (4) functionally repair tissue damage. Acellular osteochondral implants, where bone and cartilage tissue are taken together from a donor knee and processed to selectively remove cellular material, may preserve the complex architecture and molecular composition of an osteochondral allograft. Thus, decellularization and associated processing represent a straightforward and relatively simple method of acellular allograft preparation, while providing an ideal natural tissue scaffold for regeneration. Though an acellular osteochondral implant maintains the structural and composition properties of native cartilage, and may restore tissue architecture with targeted characteristics, the dense and tight-knit extracellular matrix of articular cartilage make it unclear whether recellularization is possible, and whether implants will integrate with surrounding tissue to provide long-term function and repair. This study was designed to directly test the long-term efficacy of allograft tissue, decellularized with minimal processing to retain structure and composition, and implanted into a critical-sized defect. We compared our allograft implant to an untreated defect in a contralateral joint that was repaired by drilling into the subchondral bone to create a blood clot, similar to what is commonly observed in a microfracture procedure.

We have previously evaluated acellular allografts in short-term (3-month) studies and demonstrated success in immediately providing a structural repair in a defect, but the implants showed limited integration and migration of chondrocytes into the acellular implant ^9^. However, these preliminary studies were of inadequate duration and location as the implantation and analysis was conducted in non- or low-load bearing regions of the trochlear groove at only 3 months post-surgery. Further *in vivo* investigation is needed to determine the success of a decellularized osteochondral plug in the load-bearing region of the femoral condyle, a location primarily impacted during the pathogenesis of OA. Understanding the long-term behavior and repair of a scaffold in a large animal model is paramount for predicting clinical success in humans. In this work, we aimed to measure and characterize the success of an allograft repair by investigating the structure, composition, function, and integration of an acellular osteochondral implant in a common large animal, sheep (ovine) model after 6 months *in vivo*. We prepared acellular allografts with minimal processing, removing cellular components, while maintaining important cartilage specific structural layers, mechanics, and heterogenous composition throughout the depth of the tissue. We defined the functional, structural, and mechanical characteristics of the osteochondral repair, integration zone between implant and native tissue, and the surrounding tissue using multiple analysis techniques, including displacement-encoded MRI to assess regional heterogeneity in tissue-level biomechanics. We show that acellular allografts result in a functional and integrative osteochondral repair with potential for clinical translation, but also with limitations that suggest the long-term success of the implant may suffer from a lack of cellularity and cellular migration into the repair tissue.

## II. RESULTS

### 2.1 Acellular allografts restored functional properties of native articular cartilage

To investigate the functional response of acellular allografts in load bearing regions of the femoral condyle, we analyzed whole limbs using displacement-encoded (dualMRI ^10,11^) and classical relaxation MRI with novel analysis methods to quantify tissue strain. An acellular allograft from a donor sheep was implanted into a defect in the femoral condyle on the right limb of a recipient sheep. During the surgery, we made an identical defect in the condyle contralateral (left) limb of the animals, which we left empty as an untreated control. After 6 months where animals were able to move freely, functional MRI measurements indicated that the implants dissipated loading and maintained hydration, as compared to untreated defect joints. The average strain of the implant joints under loading was significantly lower, ∼50% lower for 1^st^ principal and ∼60% lower for both 2^nd^ principal and max shear, than the strain in the untreated defect joint (**Figure 1c**). Additionally, it is known that hydration plays a vital role in the ability of articular cartilage to cushion the joint from cyclic compressive forces and to recover after loading ^12–14^. MRI relaxivity measurements of the joints before and after loading demonstrated that joints with the implant retained higher T_2_ (spin-spin relaxation) values than the untreated defect joints and the tissue that formed in the untreated defect showed the largest drop in T_2_ values from before to after loading.

**Figure 1:**
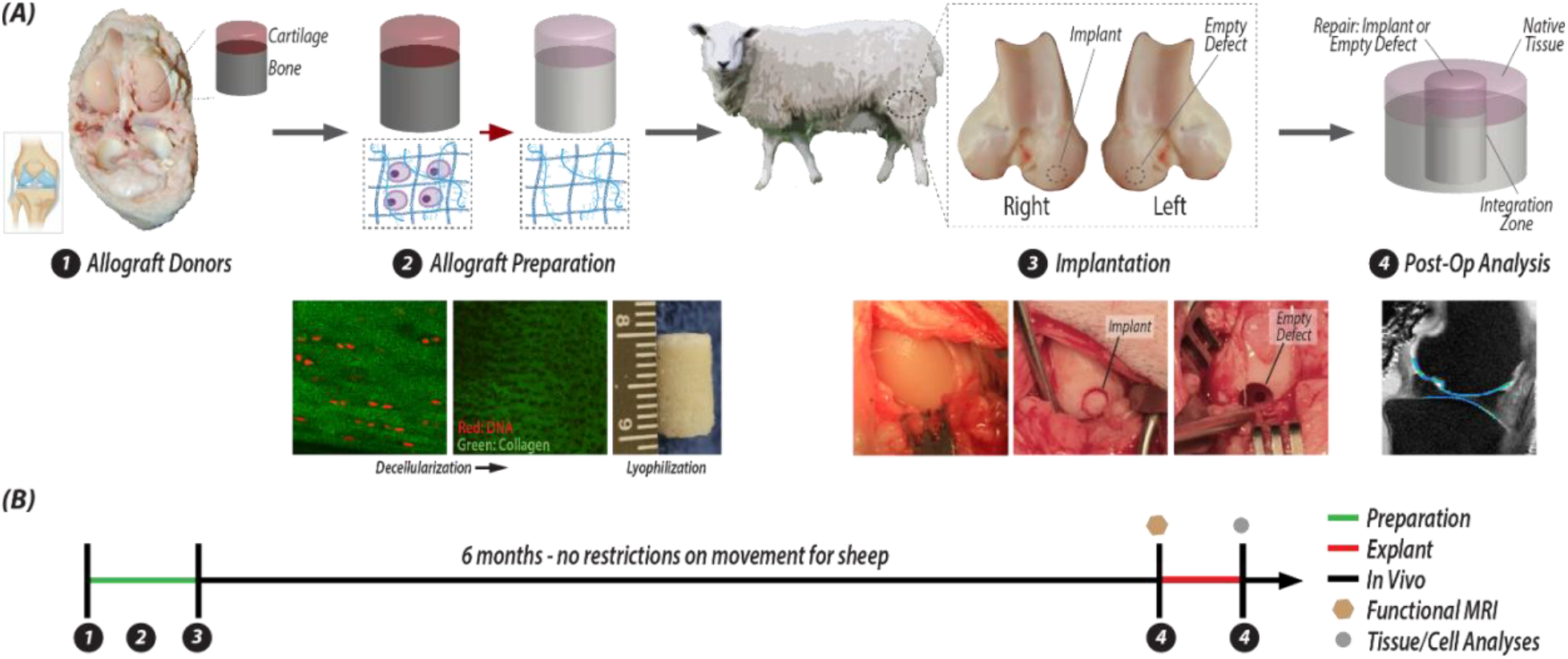
Tissue engineering strategy for integrative osteochondral repair using acellular tissues, sheep surgery, and experimental timeline. (A) Osteochondral allografts were extracted from donor sheep, and then were decellularized, lyophilized, and stored for surgery. During the surgery, the joints of receipient sheep were opened and a critical-sized defect was made in each femoral condyle. On one joint, the defect was not repaired and remained empty, and on the contralateral limb, the defect was filled with an acellular osteochondral plug. After 6 months, the repair tissue with surrounding native tissue is excised from both joints and anaylzed. (B) The timeline illustration of the steps in (A) illustrates the sequential analysis process: whole joint MRI followed by tissue- and cell-level analyses.

### 2.2 Implant architecture and structure recapitulated native tissue

To evaluate the structure and architecture of the implant and untreated defect we analyzed fixed tissue slices with histological staining and previously frozen tissue sections using Raman spectroscopy to define compositional elements of the repair. After 6 months *in vivo*, hematoxylin and eosin staining showed that implants maintained a distinct tidemark, with key division between cartilage and bone regions (**Figure 3**). When implanted tightly (press-fit, **Supplementary Figure 5**) next to native tissue, implants also maintained surface continuity from implant to native tissue. The acellular implants structurally recapitulated the thickness and typical distinct zonal layers of articular cartilage. Each cartilage zone possessed key structural differences which are essential for function, including lubrication, load support, and anchoring to subchondral bone. In contrast, the repair tissue in the empty defect failed to maintain a smooth surface with native cartilage, and instead displayed an incongruous surface even in the most successful empty joint repair (**Figure 3**). While the implants demonstrated maintenance of surface continuity with the native tissue, reduced glycosaminoglycan (GAG) content was visualized in the cartilage layer of the implant (**Figure 3**).

The implant and defect repair manifested key differences in composition. We compared the amplitude of common peaks identified from Raman spectra ^15–18^ for collagen, chondroitin sulfate, phosphate, and non-collagenous proteins on the surface of three regions in each joint: native cartilage, integration zone, and tissue (i.e., implant or defect) repair (**Figure 3**). We found consistency across all native tissue, despite whether the joint had an implant or an untreated defect. Native tissue displayed a collagen to chondroitin sulfate (CS) ratio of ∼2, a collagen to phosphate ratio of ∼14, and a collagen to non-collagenous protein ratio of ∼1.3. The implants contained a very similar composition to the native tissue: collagen:CS of 1.6, collagen:phosphate of 15, and collagen:non-collagenous protein of 2.1. Alternatively, the composition of the defect repair was different, especially in phosphate content which is an indicator of mineralization. We measured collagen:CS ratio of 1.1, collagen:phosphate of 2.9, and collagen:non-collagenous protein of 2.8, which results in differences from the native tissue by 45%, 79%, and 15%, respectively (**Figure 3**). The integration zone between the implants and native tissue in the animals retrieving the acellular implant treatment also mimicked the composition of tissue found in the animals with an untreated defect.

### 2.3 Acellular allografts restored key mechanical and tribological properties of cartilage

To determine whether the mechanical properties in the repair matched native tissue, we quantified Hertz contact and relaxation moduli using micro indentation, and measured surface tribological properties via atomic force microscopy (AFM). The implant did not demonstrate significant differences from native tissue under indentation, demonstrating a Hertz contact modulus of 393 kPa and an equilibrium modulus of 120 kPa, compared to native tissue moduli of 321 kPa and 148 kPa, respectively (p>0.05) (**Figure 4**). Defect repair sites alternatively showed a contact modulus of 414 kPa when indented, and while not significant (p<0.05), the equilibrium modulus was nearly the same as the Hertz contact modulus at 316 kPa (**Figure 4**).

The surface of native articular cartilage plays a vital role in the function of the tissue under loading and movement. Using AFM, we showed that the implant restored surface roughness and friction to values nearing that of native tissue (**Figure 4**). We measured a surface roughness on the implant of 300 nm, similar to the 321 nm roughness of native tissue, which matches previously reported roughness values of the cartilage surface ^19^. In the defect repair, we measured a significantly higher surface roughness of 500 nm (p<0.001) (**Figure 4**). Importantly, we measured a friction coefficient on the implant surface of 0.43, not significantly different from the native tissue friction coefficient on both types of joints, i.e., 0.28 in the defect joint and 0.38 in the implant joints (p>0.05). The AFM elastic modulus under indentation at the micrometer length scale (2 μm probe tip) indicated similarities between native tissue and the implant with a much softer measurement on the surface of the defect tissue (50-51 kPa on native tissue surface, 52 kPa on implant surface, 6.5 kPa on defect surface) (**Figure 4**). Friction and surface roughness properties were independently considered, as friction measured by AFM is not necessarily related to roughness of the cartilage surface and is instead influenced more by adhesion and plowing effects ^20^. Furthermore, the friction coefficient is not necessarily a good indicator of wear resistance ^21^.

### 2.4 Bone regions of acellular allograft functionally integrated with native tissue

To investigate the integration and quality of repair in the bone region of the treated and untreated defects, we used microCT imaging to quantify bone volume fraction, trabecular thickness, and trabecular spacing throughout the repair (implant or untreated defect) and integration regions. We observed integration of the bone regions with native tissue in both the implant and untreated defect joints. However, the bone in the implant repair displayed thicker trabeculae, smaller trabecular space, and increased bone volume fraction as compared to the defect joints (**Figure 5**). Furthermore, the microarchitecture of the bone in implant joints was not significantly different than the native tissue. On the contrary, in the untreated defect joints, both the bone volume fraction and trabecular thickness were significantly lower than in native tissue (**Figure 5**).

### 2.5 Implants lacked cellularity but demonstrated interfacial cellularity and integration

To evaluate cartilage integration of the repair and surrounding native tissue, we analyzed histologically stained tissue slices to visualize integration and quantify cell number in the native tissue, integration zone, and repair (implant or untreated defect). We observed some integration of the cartilage portion of the implant with native tissue, but limited cellularity within the implant. Masson Trichome staining of the repair regions (implant or defect) centered around the critical integration zone (**Figure 6**) illustrated that the integration region varied widely depending on the size of the surgical gap between the implant and native tissue. With a small gap, no new tissue filled the integration zone, but the continuity of the surface greatly improved. With a larger space between implant and native cartilage, highly cellular tissue nearly identical to the tissue in the defect repair filled the space. We quantified the cellularity in each region (i.e. native, integration zone, and repair) and confirmed that cellular density in the integration regions was nearly identical to cell density in the empty defect repair (**Figure 6**). While the tissue in the integration regions was highly cellular and seemingly fibrotic in composition, the fibrotic repair tissue succeeded in laterally integrating the implant with the native cartilage. Unfortunately, despite the high levels of cells in the integration zones, the implants did not contain many cells (**Figure 6**), illustrating an inability for cells to migrate into the implants even with a high cell number in the surrounding tissue. From previous work, we know that chondrocytes are limited in their ability to migrate much more than 100-250 μm from the surface in dense matrix, due to the limitation of their nucleus and cell size ^22^. Lack of cellularity in the implant is likely due to a myriad of factors, including high matrix density inhibiting cell migration and that the cell type in integration regions is not cartilage specific.

## III. DISCUSSION

In this work, we evaluated an acellular allograft repair as a potential clinical therapy for articular cartilage defect repair to delay the progression of cartilage tissue damage, which utilizes the advantages of autografts and allografts while reducing source limitations. Decellularized tissue has been shown to provide superior regenerative potential in many clinical settings outside of orthopedics ^23–25^, and involves minimal manipulation to produce an acellular construct that matches a tissue’s exact architecture. We showed that an acellular osteochondral implant provides cartilage-specific functional properties, and has the potential advantage that implants could be sourced from xenogenic (animal-derived) or non-viable human tissue, possibly decreasing existing source limitations.

The acellular osteochondral implants evaluated here provided force dissipation, hydration, and improved articular cartilage function as compared to an untreated defect. After 6 months, the functional dualMRI data indicated that the osteochondral implants had physically integrated laterally into the surrounding native tissue as the implants were able to transfer strain throughout the joint, rather than concentrating the strain near cartilage damage, as seen in the dualMRI performed on untreated defects. These indicators of improved function after six months in a load-bearing region were promising and motivated us to look at the repair and integration of the implant in detail. MRI relaxivity data also demonstrated that the acellular implant and native tissue T_2_ values were not significantly different, while the tissue in the untreated defect had significantly lower T_2_ values (**Figure 2)**. Since T_2_ can be used as a surrogate of tissue water content ^26^, these data suggest that the implant preserved or restored critical hydration. Additionally, the architecture measurements taken with microCT of the bone regions of the implants (**Figure 5)** further suggest that the tissue was experiencing loading, which in turn signaled to bone cells to remodel and strengthen trabeculae. The microCT data of bone structure supports our conclusion from MRI results that the implant experienced loading and successfully dissipated the force. Histological assessment further supports the functional data, illustrating that the cartilage regions of the untreated defect repair were recessed from the surface (**Figure 3)**. As a result, the tissue in the untreated defect would have experienced less direct loading, leading to the thinner and more widely spaced bony trabeculae and lower bone volume fraction, as compared to implant and native tissue.

**Figure 2.**
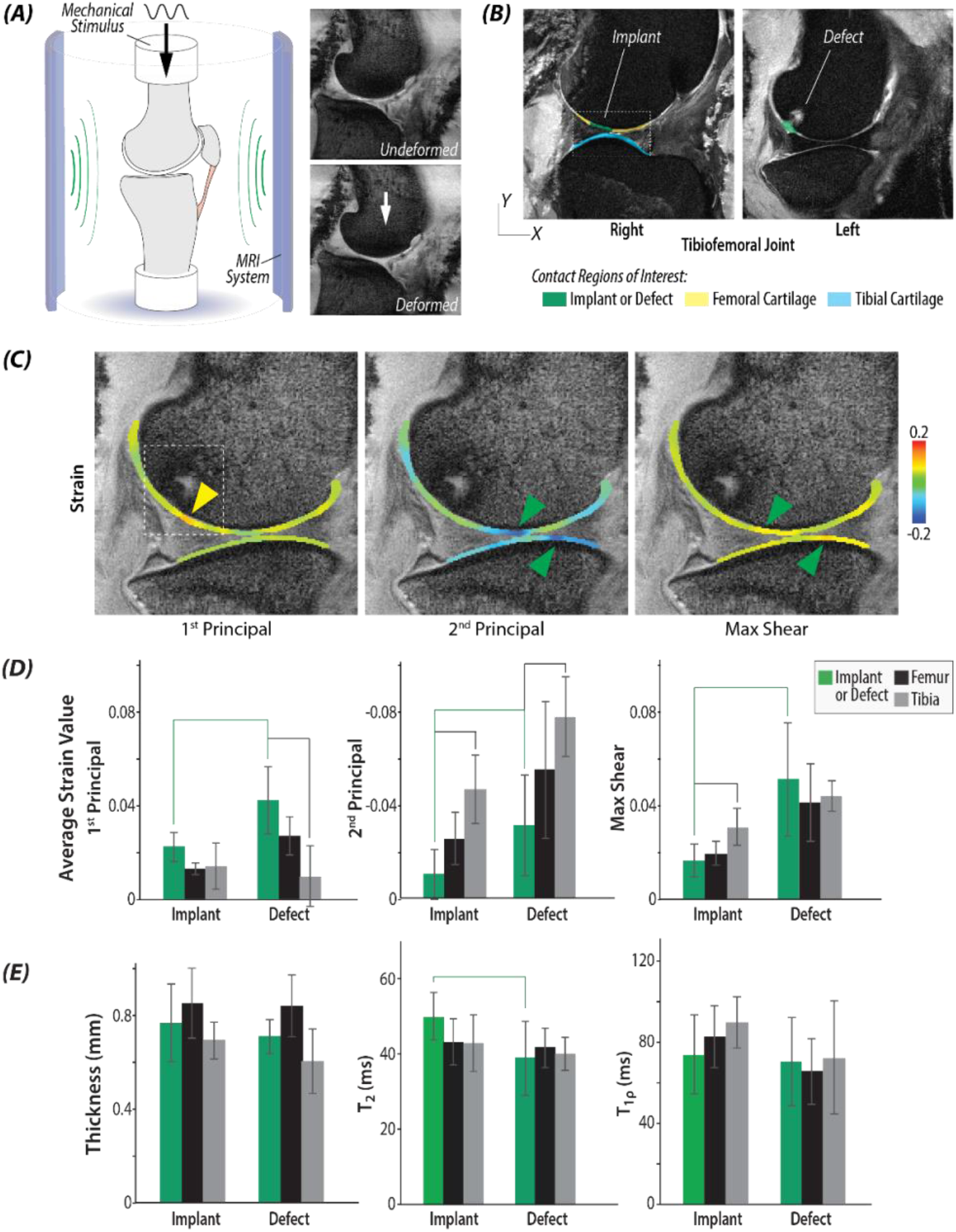
MRI whole-limb functional analyses indicate mechanical and structural similarities between the implant and native cartilage. (A) Schematic setup of the experiments with morphologic images of the tibiofemoral joint in undeformed and deformed positions, with load magnitudes and timing mimicking the gait cycle. (B) Morphological images show positions of the implant and defect, with respect to native, surrounding cartilage. (C) Principal strains were calculated using dualMRI-based phase contrast mapping during cyclic loading. (D) Average strain value of principal strains and max shear strains in the region of implant, defect, femur and tibia. Green brackets represent significant differences between implants and defects (p<0.05). Black brackets represented significant difference between implants/defects with other regions on femur or tibia (p<0.05). (E) Thickness of the implant or defect was not measurably different from surrounding cartilage. Relaxometric analysis indicates biochemical differences between the the implant and defect, as evidenced by T_2_, but not T_1_*ρ*, mapping. The green bracket indicates marginally significant difference between implant and defect (p=0.079). (n=6 defect joints and 6 implant joints).

**Figure 3.**
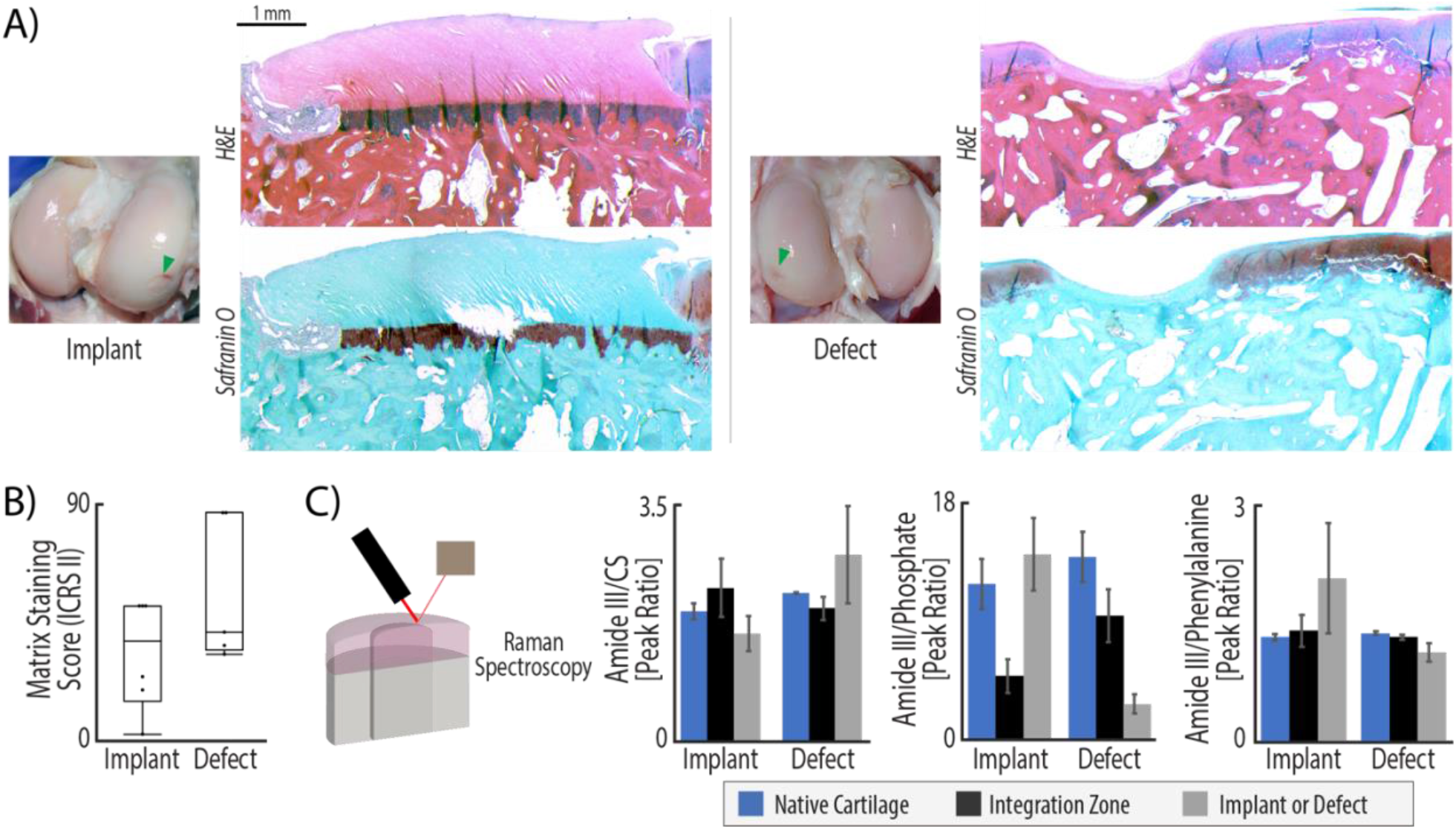
Structure and composition of implants closely matched native tissue. (A) H&E staining illustrates that the implant fills the injury void and maintains both cartilage structure and surface continuity with native tissue. In contrast, defect repair produces a much thinner layer of repair and lacks surface continuity. Safranin-O/fast green staining demonstrated that acellular allografts fail to maintain many glycosaminoglycans (GAGs), which is a known consequence of decellularization (GAGs stain red). (B) ICRS II scoring of metachromasia, or matrix staining, was evaluated in safranin-O and fast green tissue stained slices. Full metachromasia (a score of 100) would be native tissue distribution of glycosaminoglycans (GAGs). Both repair types display matrix staining scores of ∼50, illustrating some GAG staining in the repair, but far from native levels. (C) Raman spectroscopy quantifies compositional differences between the two types of repair. The collagen to chondroitin sulfate (Amide III/CS) ratio indicates higher levels of relative collagen in the defect repair and integration zones, while native and implant tissue maintain similar ratios of collagen to chondroitin sulfate. The increase of collagen in the defect repair suggest that these regions are repaired with fibrotic tissue. Additionally, defect repair and integration regions are highly mineralized compared to native tissue and implant repair (ratio of 3 vs. a ratio of 15), indicated by the Amide III/Phosphate ratio. While not statistically significant (p<0.05) due to the high levels of variance in samples of each group, the trends indicate that implant tissue structure and composition matches native tissue. (n=6, statistics run with a two-way ANOVA test to determine cofactor significance, followed by a Tukey’s honest significant difference test to determine p-values between implant types or tissue zones).

Surface lubrication and low friction are additional key components to a successfully engineered tissue construct for cartilage repair. Cartilage injury in OA leads to a loss of imperative surface lubrication that facilitates smooth sliding during joint movement ^27^. Surface measurements using atomic force microscopy revealed that the acellular osteochondral implant restored surface roughness and friction values to those measured in the native tissue of the sheep (**Figure 4)**. Alternatively, the empty repair (untreated defect) formed a significantly rougher surface, suggesting the possibility of altered contact during sliding that may lead to increased wear of the cartilage counter face – a hallmark of further degradation in the joint. In contrast to the composition of acellular implants, untreated defects filled with non-cartilaginous, mineralized tissue. The Raman spectra analysis of tissue composition in each repair condition (implant or empty/untreated) compared to native tissue indicate that defects left untreated filled with a tissue that is high in collagen I (45% increase from native tissue) and mineralized. The change in collagen to phosphate ratio from 14-15 in the implant and native tissue to 2.9 in defect repair (79% increase in phosphate content) is substantial (**Figure 3)**. High quantities of phosphate lead chondrocytes towards apoptosis, which is the first step in endochondral ossification and bone formation ^28,29^. The bulk mechanical profile of the defect repair supports the Raman spectral analysis of composition positing that the defect repair may be fibrotic and high in mineral content, contributing to limited tissue relaxation after loading (**Figure 4**). Unfortunately, the integration region between the implant and native tissue was composed of tissue with the same properties (composition, histology, mechanics, and surface tribology) as the tissue measured in the untreated empty defects. This parallel composition of the defect and integration zone is consistent with the idea that the gaps between implants and native tissue fill with the same blood and bone marrow components post-injury as that observed in the defect.

**Figure 4:**
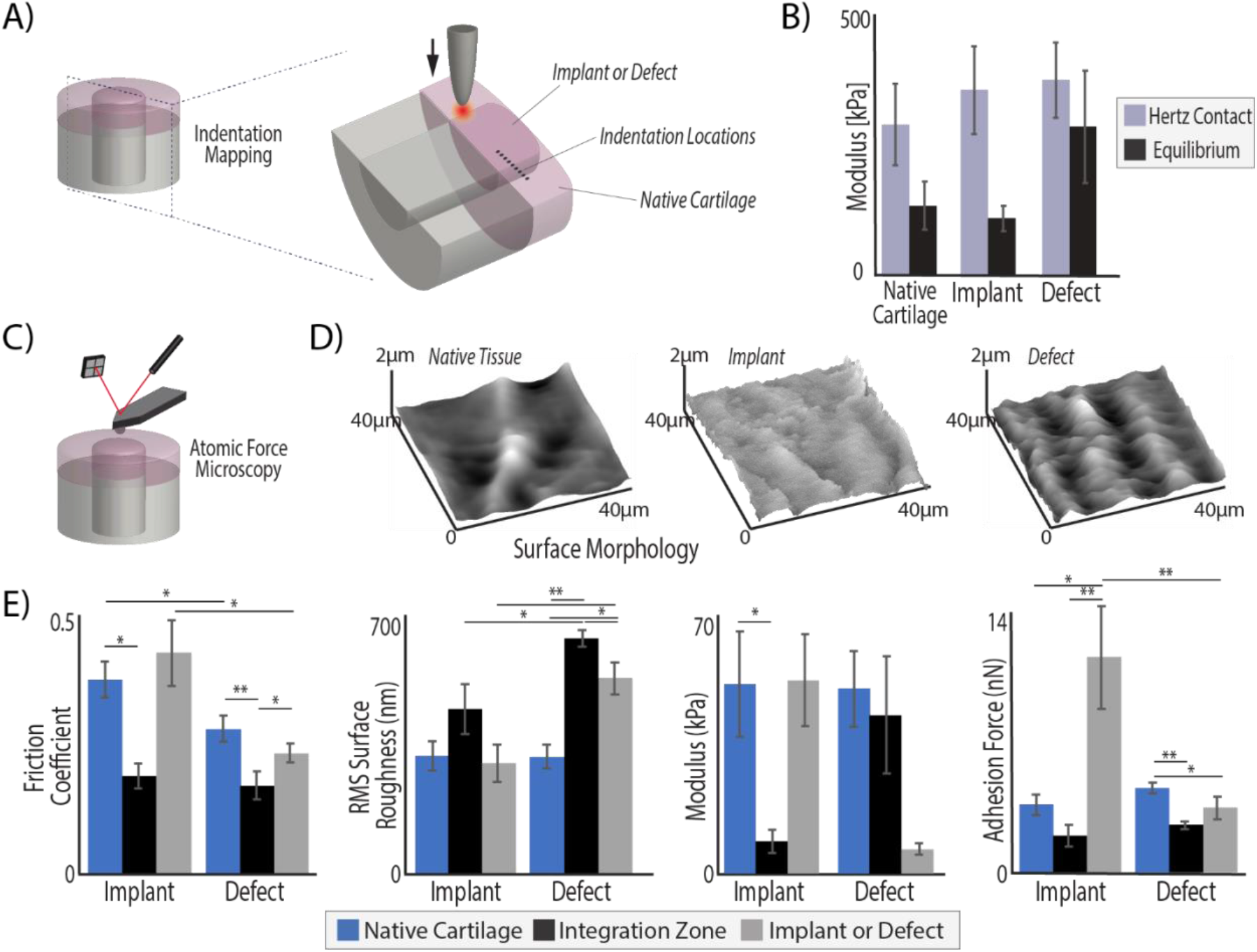
Acellular allografts restore native cartilage mechanical and surface properties. Acellular osteochondral implants, but not untreated defects, achieved mechanical and tribological surface properties of surrounding native tissue regions. (A) Bulk tissue analysis was performed using micro indentation (100 μm tip) in the middle zone of cartilage on all explanted tissues at nine, evenly spaced (200 μm apart) locations, centered around the integration zone. (B) The Hertz contact modulus under indent compression was consistent among the different implants and native tissue, but the empty defect tissue fails to demonstrate much relaxation, where the implant behaved identically to native tissue (equilibrium moduli= ∼100 kPa). The measurements from all four points in each tissue region were averaged, resulting in a representative value for the whole tissue region but leading to increased standard deviations. (C) Atomic force microscopy (AFM) was performed with a 2 μm spherical probe to (D) map the surface of each analysis region and (E) quantify frictional coefficients, surface roughness, micro moduli, and adhesion force. The implant tissue restored native values of friction, surface roughness, and micro moduli. In contrast, the surface roughness and modulus of defect repair and integration regions were significantly different than native tissue. (*p < 0.05, **p < 0.001, n=6, statistics run with a two-way ANOVA test to determine cofactor significance, followed by a Tukey’s honest significant difference test to determine p-values between implant types or tissue zones).

**Figure 5:**
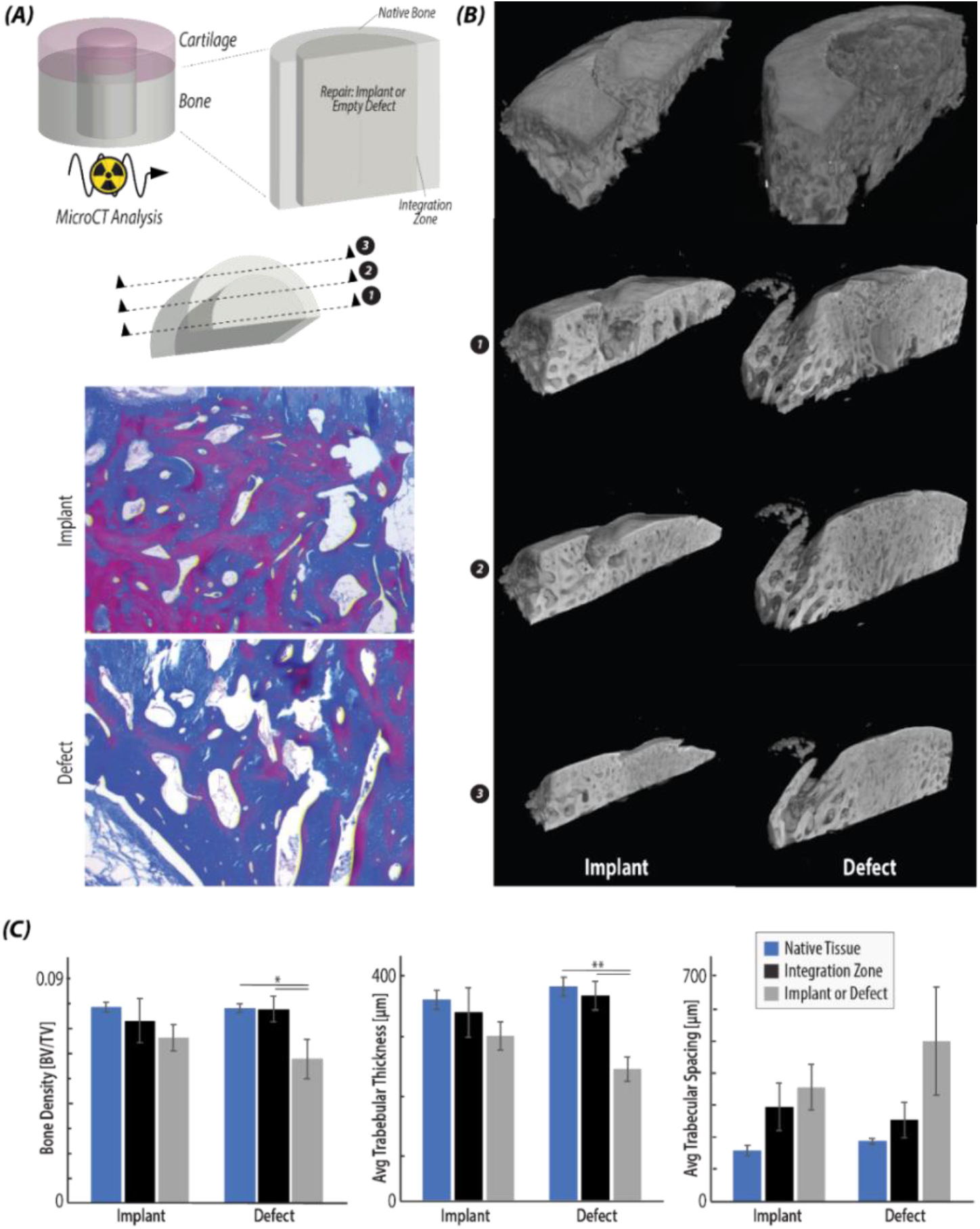
Acellular allografts demonstrate bony integration with native tissue. (A) Radiographic (μCT) analysis was used to visualize and quantify bone regions of the native tissue, integration zone, and repair tissue (i.e., implant or defect). Masson’s Trichrome histological stain was used to visualize 2D regions of the bone in each type of repair (implant/defect) (B) Reconstructed μCT scans of each type of repair show that the bone regions of the implant repair maintain surface continuity and a subchondral bone layer, while the defect tissue does not contain the thin upper layer of subchondral bone. Reconstructed slices from the bisected plane towards the outside of the implant/defect confirms the histological images in (A), as the trabeculae are thicker in the implant repair than the defect. (C) Quantification of the bone volume/tissue volume, the average trabecular thickness, and the average trabecular spacing in each of the regions of interest confirms that the defect repair contains a significantly decreased bone tissue volume and trabecular thickness, while the implant repair is not significantly different from native tissue. (*p < 0.05, **p < 0.001, n=6, statistics run with a two-way ANOVA test to determine cofactor significance, followed by a Tukey’s honest significant difference test to determine p-values between implant types or tissue zones).

Several important recent findings posit that integration with native tissue is critical to the long-term success of any engineered cartilage tissue. Furthermore, many researchers have shown that it is essential for an implant to promote cellularity in order to successfully integrate with native tissue ^2,30^. In articular cartilage, promoting cellularity is a distinct challenge as the dense extracellular matrix limits cell movement ^22^, the tissue is non-vascular, and even native tissue displays a low cell density compared to many tissues. While our work demonstrates acellular allografts provide a structural and functional repair under loading similar to native tissue, integration with bone, and some physical lateral cartilage integration, we also show that even after 6 months *in vivo*, implants failed to promote cellularity within the implant cartilage despite high levels of cellularity in the integration zone, which is potentially a significant drawback to the repair method. We believe that targeted future work focusing on this limitation could advance the clinical effectiveness of acellular osteochondral implants. Coating the implants or surgical site prior to implantation with cartilage specific growth factors such as IGF-1 has been shown to stimulate cartilage specific cell migration to the border regions ^31,32^. While we observed high cellularity in the integration region of the defect repair (**Figure 6**), the cells were likely primarily of a fibroblast phenotype that failed to migrate into the acellular cartilage implant. Combining cartilage-specific growth factor treatment with an acellular allograft could be an interesting extension of the repair strategy explored here, with potential for improved integration and effectiveness of the repair. Furthermore, it is evident from the results presented here that the surgical technique and careful implantation also affects the success of the implant. In animals where the implant was tightly press-fit within native tissue, and the surface of the implant and native tissue were aligned, we observed improved surface continuity and functionality (**Figure 3, 6, and Supplementary Figure 5**). Prioritizing specific and precise implantation techniques would result in a decrease in the volume of repair (e.g. fibrous) tissue in the integration region as well as superior surface continuity between the implant and native cartilage. Finally, recent work posits that while the dense cartilage extracellular matrix limits cell migration into an acellular allograft, breaking down the tissue into particles facilitates cell migration while maintaining cartilage specific structure and mechanics ^22,33,34^. Future work investigating an implant where the cartilage portion of the osteochondral implant is first pulverized could improve lateral integration, cellularity, and clinical effectiveness ^22^. However, it is not clear whether the use of pulverized and reconstituted cartilage would potentially lose the benefit of closely mimicking the tissue structure and function that were key benefits of the acellular implant used herein. Future studies may need to balance structure, function, and cellularity to improve integrative cartilage repair.

**Figure 6.**
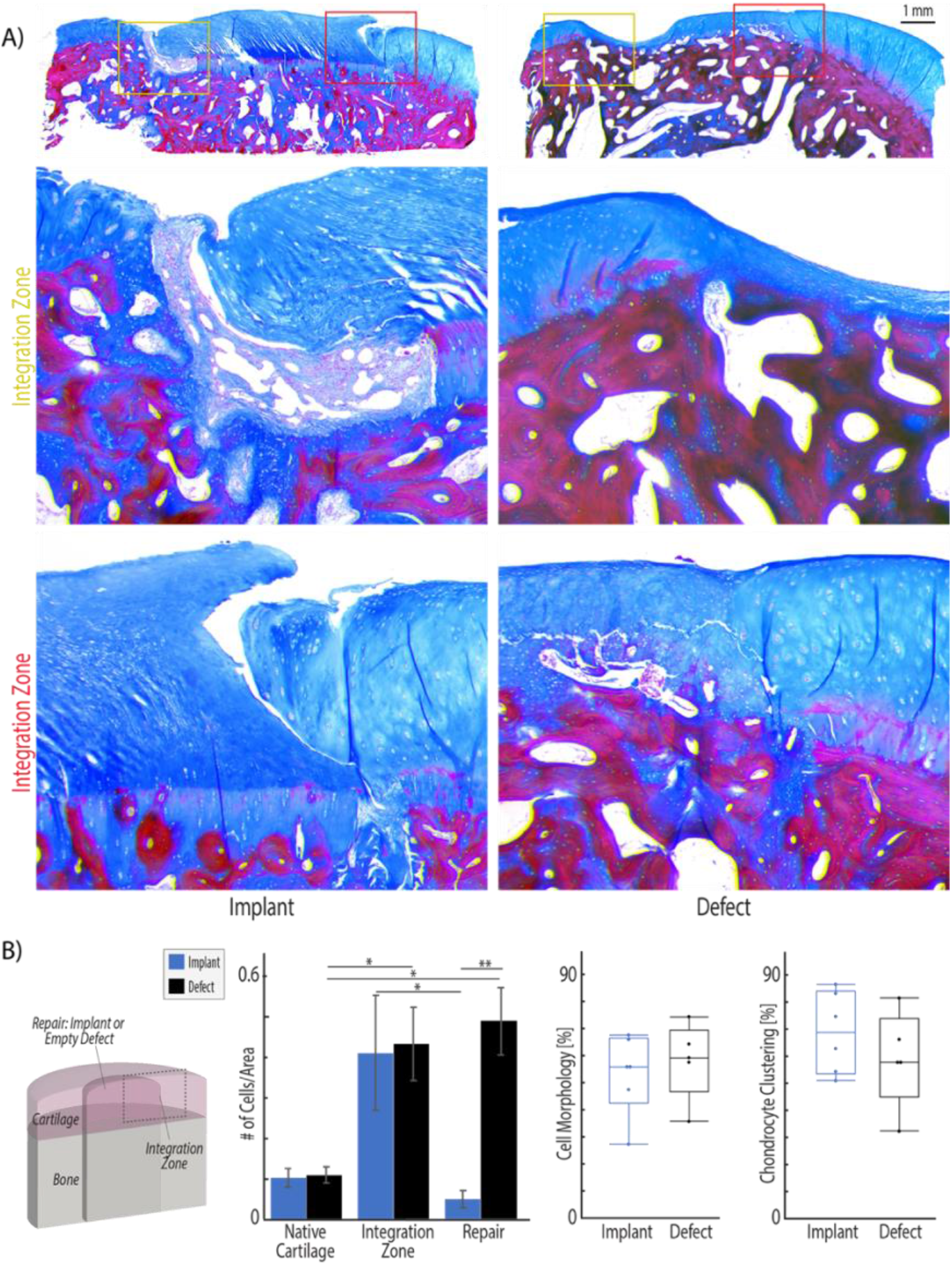
Acellular allografts promote high cellularity in the integration zone with native tissue. (A) After 6 months *in vivo*, Masson’s trichome staining shows that integration regions of implants are either filled with tissue which is densely packed with cells and appears fibrotic (top image) or no new tissue forms when the gap between implant and native is small (bottom image). The integration region of defect repair tissue illustrates a transition from native cartilage with sparse, evenly distributed chondrocytes to a fibrotic tissue with visible collagen fibers and densely packed cells. (B) Masson’s Trichrome images were analyzed using ImageJ thresholding and particle counting to quantify cell density in each region. The integration regions and defect repair tissue demonstrate significantly higher cellularity than native cartilage tissue. Additionally, while the integration regions are highly cellular, the cells do not migrate into the implant. Finally, ICRS II scoring was used to evaluate cell morphology. Unhealthy chondrocyte clustering is increased in the defect samples and can be observed near the integration zone in the histology examples in (A). Additional ICRS II tissue metrics are reported in supplementary figure 4. (*p < 0.05, **p < 0.001, n=6 animals, statistics run with a two-way ANOVA test to determine cofactor significance, followed by a Tukey’s honest significant difference test to determine p-values between implant types or tissue zones).

Long-term function of defect repair strategies in a relevant, non-human model is a critical aspect to any clinical intervention or engineered tissue construct prior to human clinical trials. Large animal models (e.g. in sheep, goat, dog, and horse) are useful in analyzing the effectiveness of a treatment and closing the gap between *in vitro* experiments and human studies ^35,36^. Sheep joints specifically demonstrate joint size, cartilage thickness, and limited intrinsic healing ability amenable to human joints ^37^. As a result, many researchers utilize a sheep model ^36^ to show tibiofemoral defect treatment efficacy and limitations of microfracture with additional treatments ^38–40^, modified autologous plug transfer procedures ^41–45^, variations of allografts ^9,46,47^, and tissue engineered scaffolds ^48,49^. While all studies in load bearing regions of a sheep elucidate the potential clinical success of a technique or scaffold, many referenced above are only 1-3 months long, and thus do not indicate the long-term success or degradation. Additionally, while powerful tools, sheep models are limited to answering specific questions. Unlike humans and dogs, it is difficult to manage post-operative non-load bearing or therapy sessions in sheep. Therefore, sheep are best used to evaluate repairs for which post-operative management is minimal, and animals can bear weight immediately ^37^.

## IV. CONCLUSION

The work presented here shows that after 6 months in a sheep knee, implants match the structural composition of surrounding native cartilage, provide bulk mechanical properties under compression, enable the critical capability of load transfer throughout the joint, and restore surface roughness and friction. Importantly, here we show that acellular osteochondral implants are a viable and potential strategy to postpone osteoarthritis progression and patient pain in joints by restoring function, for months to years. However, the limitations of cellularity in the repair could limit the longevity of the effectiveness of this repair. Additionally, the analysis suite presented here used to evaluate repair from the whole joint to the cellular level provides a model for evaluating new cartilage repair strategies, including novel and non-invasive dualMRI analyses to incorporate strain measurements into osteoarthritis diagnosis and treatment evaluation. Using these analysis tools, we have demonstrated effectiveness of a surgical intervention that requires only one surgery, does not rely on a limited tissue source, can immediately restore articular cartilage structure and function, and maintains cartilage-specific function for at least 6 months in a relevant animal model. For the more than 50% percent of patients with osteoarthritis in their knee who are younger than 65 ^3^, acellular osteochondral implants could provide a potential solution to decrease pain and increase function until the age where a total joint replacement is recommended.

## V. METHODS

### 5.1 Decellularization of Osteochondral Allograft Implants

Ovine osteochondral implants were harvested from the load bearing region of the distal femur of (donor) stifle joints using the Osteochondral Allograft Transfer System (6mm diameter; OATS, Arthrex, Naples, FL). Implants were decellularized in 2% Sodium Dodecyl Sulfate (SDS, Sigma Aldrich) at 37°C under constant agitation for 8 hours, washed in phosphate-buffered saline (PBS) and then incubated with 3.3 mg/ml DNase, 50 mg/ml RNase, 1% P/S, 1% fungizone at 37 °C under agitation for 24 hours. To deactivate the DNase after treatment, implants were bathed in 0.02% EDTA, with a final PBS wash. Acellular allografts were flash frozen in liquid nitrogen and lyophilized for 48 hours prior to surgical implantation.

### 5.2 Surgical Procedures for Cartilage Defect Repair Using Decellularized Allografts

Cartilage defect repair using acellular allografts was performed on six ewes aged 2.25 ± 0.43 years, who all fell under a similar weight category (74.8 ± 8.4 kg), as approved by the Purdue University Institutional Care and Use Committee. Using an end mill system, osteochondral defects were placed via drilled holes in the load bearing region of both medial condyles (6mm diameter and 10mm depth). An acellular allograft was press fit into the defect region on the right stifle joint, and the identical defect on the contralateral left joint was left empty as an untreated control. The joints were closed in layers, and the animal was allowed to move freely with no weight bearing limitations post-surgery. After six months, ewes were sacrificed, and stifle joints were frozen at -80°C until subsequent analysis. Following MRI analysis (discussed below), repair tissue was harvested by exposing the stifle joint space and removing tissue with a scalpel. Implants and surrounding native tissue (10 mm diameter, 10 mm depth) were extracted from each joint and bisected, with one half stored at -80°C and the other half fixed for 48 hours in 4% paraformaldehyde (PFA).

### 5.3 Whole-Joint Analysis of Structure and Mechanical Function by MRI

Whole explanted joints were secured with polymethylmethacrylate into a custom-machined loading apparatus. The loading device fixed the tibia while allowing the femur to move in compression-distraction under mechanical stimulation, and each specimen was secured with a natural flexion angle of 135°. Joints were scanned in a 7.0 T MRI scanner (Bruker Medical GmbH, Ettlingen, Germany) for morphology, relaxometry, and strain mapping. For morphology imaging, Rapid Acquisition with Refocused Echoes (RARE) and Fast Low Angle Shot (FLASH) pulse sequences were applied in sagittal plane to cover the whole joint. RARE imaging parameters were: TE/TR=22.88/3000 *ms*; NA=4; in plane resolution=0.25×0.25 *mm*^*2*^; image matrix size = 256×256 pixels^2^; slice thickness = 2.0 mm; RARE factor=8; number of slices=25. FLASH imaging parameters were identical, with the following modifications: TE/TR=4.806/390.2 *ms*; NA=4; flip angle = 30°. Thickness measurements of the full thickness of cartilage were obtained from morphology imaging using manual measurements (Fiji).

For relaxometry, the Multi Slice Multi Echo (MSME) sequence was used for T_2_ (spin-spin relaxation) measurements, with the following imaging parameters: TE/TR=7.781/3000 *ms*; number of slices=25. The RARE sequence was used for T_1_*ρ* relaxation measurements, with the following imaging parameters: TE/TR=34.96/2578 *ms*; NA=2; Repetition=5; Echo spacing=8.74 *ms*; RARE factor =8; spin lock strength=851 *Hz*; spin lock duration=10, 45, 80, 115 *ms*. Relaxation measurements were obtained using monoexponential fitting routines of the time-course data.

For strain mapping, we utilized dualMRI ^11,50,51^, the stimulus was applied at the femur, moving the joint from undeformed to deformed. The DENSE-FISP sequences were used to scan the joints with an *xy* displacement encoding gradient area of 0.507 *π*/*mm*. The scan was synchronized with cyclic loading at frequency of 0.33 Hz with 445 *N* compression and 100 *N* retraction. With the same slice resolutions and dimensions introduced before, FISP sequence has parameters of TE/TR = 2.505/5.01 *ms*; NA = 4; number of segmentation = 2. To study the effects of the external loading on the different repair types (i.e., implant or defect), analysis focused on the loaded cartilage images. The dual MRI loading setup was designed to align the implant or defect at the direct compression point, though they might not be located exactly at the cartilage-cartilage contact region during image. The displacement was smoothed using locally weighted linear regression smoothing (LOWESS) in MATLAB (MathWorks, Natick, MA). To avoid variation due to joint orientation, coordinate independent strains were calculated.

### 5.4 Structural Analysis Using Histology

Matrix deposition, glycosaminoglycan content, and overall repair structural continuity and quality were evaluated using histology. One half of the extracted tissue from each joint was fixed for 48 hours in 4% paraformaldehyde, decalcified for 3 weeks in EDTA, dehydrated progressively with EtOH, embedded in paraffin, sliced into 5 μm sections using a microtome, and mounted on microscope slides. Sections were stained individually with Hematoxylin and Eosin (H&E) and Safranin-O/Fast Green (Histology Core, CU Denver Cancer Center). The International Cartilage Repair Society II (ICRS II) scoring paradigm was used to evaluate relative tissue quality in each repair (i.e. implant or defect) ^52^. Scoring was performed by three independent and blinded observers.

### 5.5 Structural Analysis Using Raman Spectroscopy

The composition of tissue regions was quantitatively evaluated using Raman spectroscopy with a 785nm laser and spectra collected for wavenumbers 400-1600 cm^−1^. Raman spectroscopy data was collected on an upright inVia microscope (Renishaw, Wotton-under-Edge, UK). The regions measured previously in atomic force microscopy were first identified using brightfield microscopy at 5× magnification, assisted by India ink markings on the edge of each analysis location. Samples were submerged and a 63× immersion objective was focused with a 785 nm laser to illuminate a ∼1 μm diameter spot on the surface. In each region, spectral maps consisting of sixteen measurements covering a 40 μm x 40 μm area were collected. At each point in the spectral map, cosmic rays were removed from the collected spectra, a linear baseline was subtracted and intensity normalized, and spectra were smoothed ^18^. The sixteen spectral acquisitions that composed each spectral map were averaged, key peaks of interest were identified, and peak amplitude was calculated using custom R code following a previously established Raman spectra analysis approach ^53^. Relative peak amplitudes for peaks of interest (amide III-1280 cm-1, *ν*_2_ phosphate-441 cm-1, chondroitin sulfate-1068 cm-1, and phenylalanine-1003 cm-1) ^15–17^ were compared to calculate relative compositional differences in each region.

### 5.6 Characterization of Bulk Mechanical Properties via Indentation

To assess the *ex vivo* bulk mechanical properties of the native cartilage, integration zone, and tissue (i.e., implant or defect) repair, microindentation was performed at nine points spanning 1.6 mm (200 μm spacing between points) centered at the integration zone (**Figure 4**). In preparation for microindentation a flat, 1-2mm section of tissue was sliced; the resulting test surface contained a cross-section of repair tissue, integration region, and native cartilage. Indentations (Hysitron TI 950 TriboIndenter, xZ-500 extended displacement stage) were performed on submerged samples using a 100 μm radius spherical probe. For each indent, the probe was first lifted off the sample surface, then indented to a depth of 20 μm at a rate of 240 μm/s, the indent was held for 45 seconds to permit force relaxation. A virtual contact point for each indent was determined following established methods ^54^. Hertzian contact modulus (assuming ***ν***= 0.5) and equilibrium modulus (assuming ***ν***= 0.0 after fluid drainage) were calculated at each indent site. To calculate the reported regional bulk mechanical properties, the moduli at each of the four points within each region were averaged.

### 5.7 Surface Property Characterization by Atomic Force Microscopy (AFM)

Articular cartilage surface roughness, frictional coefficient, adhesion, compressive modulus, and topographical structures were measured using atomic force microscopy (Keysight 5500 AFM, Keysight Technologies Inc., Santa Rosa, CA, USA). All measurements were taken on the articular surface of the implant in the three key regions of interest: native cartilage, integration zone, and tissue (i.e., implant or defect) repair. A cantilever with known geometry was used (2 μm borosilicate sphere, NovaScan), and the cantilever stiffness was pre-calibrated to 0.36 N/m by the thermal fluctuation method ^55^. Lateral calibration was determined using an improved wedge calibration method ^56^ and a TGF11 silicon calibration grating (Mikromasch). Each sample was affixed to a μ-Dish high grid-50 dish (Ibidi, USA) using a viscous cyanoacrylate and hydrated for all AFM testing. At each location, the macroscopic region of interest on the articular cartilage surface was approached and a brief scan (4×4) was performed to ensure the height change was in an acceptable range (∼7 μm). After locating a measurable 40×40 μm area, sixteen independent locations of the scan area (4×4) were measured in force-volume mode, indented at 5 μm/s with a setpoint force of approximately 12 nN. To estimate the compressive modulus, force-displacement approach curves were fit to the Hertzian linear elastic model for a defined round tip geometry ^57,58^. Adhesion force was measured as the pull-off force upon tip separation from the surface during probe retraction. Both adhesion and compressive modulus were extracted from force-distance curves using PicoView 1.14 AFM analysis software. Immediately following and within the same scan area, a high resolution contact scan (1028×1028 px, or 40nm/px) was performed at 20 μm/s with a constant applied normal force of 20 nN to generate detailed topography, raw deflection, and lateral voltage trace/retrace signals of surface features. Areas where the controller overloaded the sample or did not interact with the surface were excluded. Topographical images were analyzed to calculate root-mean-square (RMS) surface roughness using Gwyddion 2.56 SPM analysis software ^59^. The averaged lateral voltage signal over each area was converted to friction force by multiplying by the lateral calibration constant and dividing by the normal force to yield the coefficient of friction.

### 5.8 Quantitative Imaging of Osteochondral Bone Repair and Remodeling

Trabecular bone volume fraction, trabecular thickness, and trabecular spacing were measured using micro-computed tomography (XRadia Versa 520, Zeiss, Dublin, CA, USA). The explants were scanned with a 0.4× objective, voxel size of 13.4 μm^3^, energy settings of 50 kV, 3.0 W, 3 s exposure, 801 projections, and using the LE3 or LE4 filters. Automated centering and beam hardening corrections were applied using Scout-and-Scan Control System Reconstructor software (v 14.0.14829.38124). To analyze the bone structure, collected micro tomography images were imported into Dragonfly software (ORS v 4.1) and the Otsu algorithm ^60^ was implemented to separate ranges of the histogram corresponding to bone and background noise. Distinct volumes to analyze were designated in three key regions: native cartilage, integration zone, and tissue (i.e., implant or defect) repair. In each region, trabecular bone volume fraction was calculated. A sphere fitting-method was used in each trabeculae or in the space between trabecula to calculate trabecular bone volume fraction, trabecular bone thickness, and trabecular bone spacing using Dragonfly Bone Analysis.

### 5.9 Cell Quantification using Histological Images

Cellularity of all regions of interest and cell migration into the repair tissue was quantified in each of the three key regions using paraffin embedded tissue slices (preparation detailed above) stained with Masson Trichrome (Newcomer Supply) following the manufacturer recommended staining protocol after tissue hydration with decreasing concentrations of EtOH. Each region (native, integration zone, and tissue repair) was isolated in a high resolution image and standard ImageJ thresholding ^61^ was used to identify cells (blue/black spots) and quantify cell number/tissue area using Image J particle counting.

### 5.10 Statistical Analysis

To test our hypotheses that treatment (osteochondral implant or empty defect) influenced articular cartilage architecture and repair (stiffness, surface roughness, adhesion, protein signature, bone composition) *in vivo*, mixed model analyses of variance (ANOVAs) were performed on generalized linear models with treatment, animal, and implant location (left stifle joint or right stifle joint) as the predictors, and the resulting measurement variable as the response. In all data sets, if treatment, animal, or implant location was found to be a significant predictor then the p-value significances between each group was calculated using Tukey’s Honest Significant Difference Test. The same statistical approach was used for data collected in each region of interest. To compare between regions with the same treatment, the ANOVA was performed on a generalized linear model with region, animal, and repair type as predictors. Statistical significance in all experiments was defined as p<0.05. All statistical testing was performed using R software and the lmer package.

## VI. ACKNOWLEDGEMENTS

The authors acknowledge the histology core at the University of Colorado at Anschutz Biorepository Core Facility for technical assistance with embedding, sectioning, and H&E staining. Financial support is gratefully acknowledged from NIH R01 AR063712, DoD / CDMRP W81XWH-20-1-0268, and the Purdue University OVPR Emerging Research Incentive Grant Program.

## VII. AUTHOR CONTRIBUTIONS

Conceptualization, L.C, J.B. and C.P.N.; Methodology, J.B., L.C, K.P.M., K.F, K.E., V.F, and C.P.N.; Animal Surgery, G.B.; Formal Analysis, J.B., L.C., K.P.M., K.E; Writing – Original Draft, J.B., Writing – Review & Editing, All authors; Statistical Analysis, J.B, N.E.; Funding Acquisition, C.P.N.; Resources, G.B., V.F. and C.P.N.; Supervision, C.P.N.

## SUPPLEMENTAL INFORMATION

**Supplemental Figure 1.**
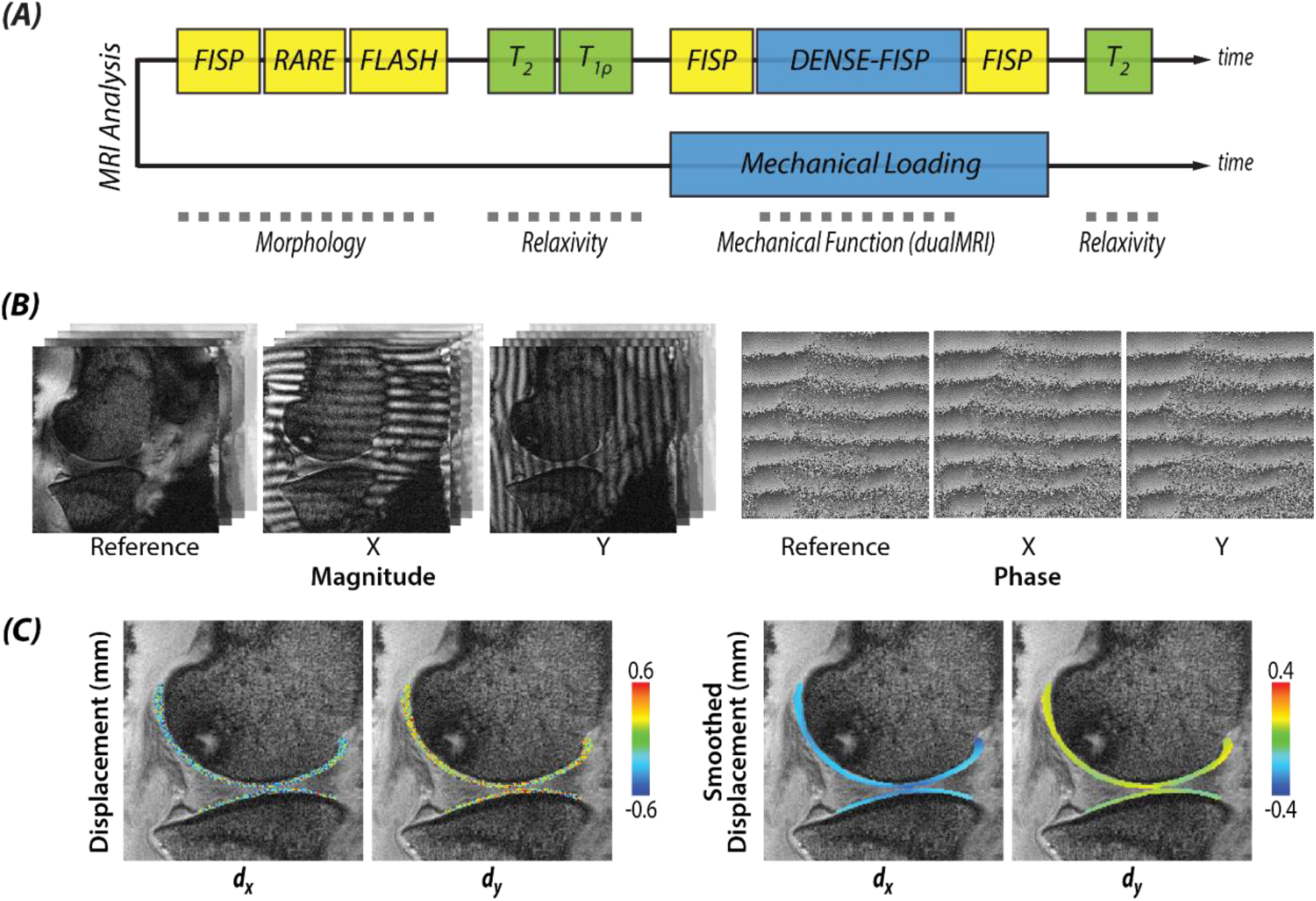
MRI Methods. (A) After 6 months, sheep were necropsied and evaluated with MRI including morphology, quantitative and displacement-encoded scanning. (B) Measured magnitude and phase data required for evaluation of intratissue mechanics during cyclic loading. The (C) displacements were calculated to determine resulting strain measurements.

**Supplemental Figure 2.**
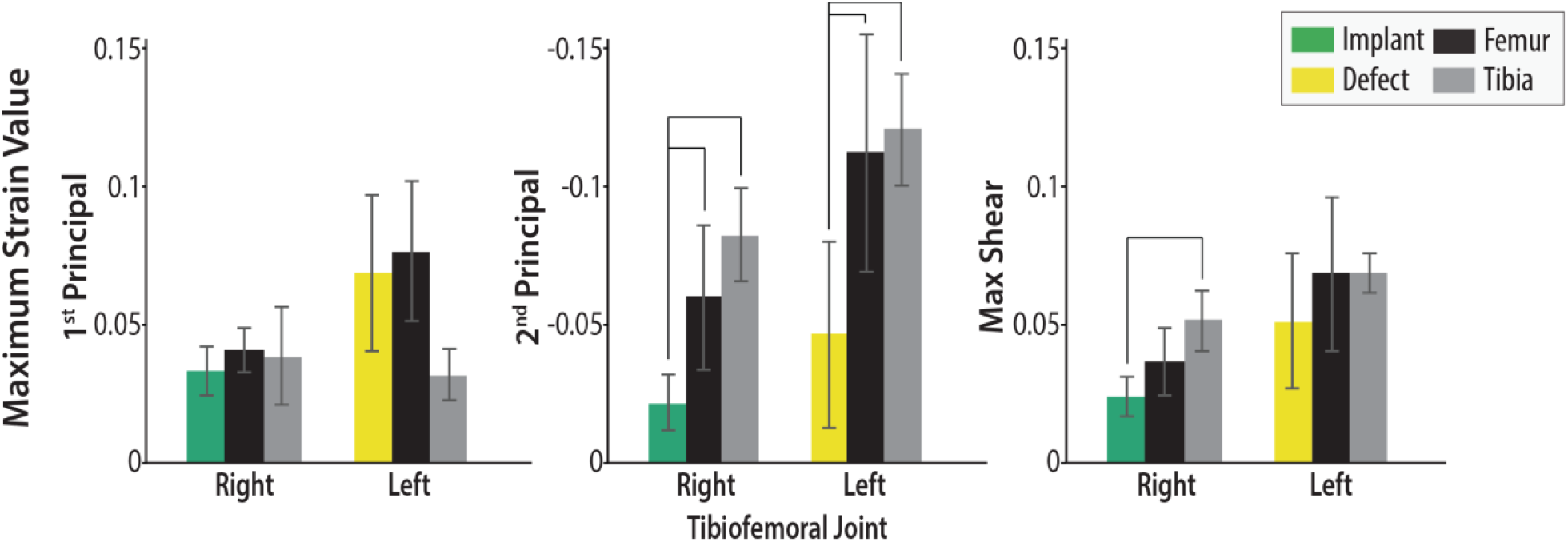
MRI Max Strain. Maximum strain values in the implant, defect, femur, and tibia mimic the same patterns of the average strain values reported in Figure 2.Black brackets represented significant difference between implants/defects with other regions on femur or tibia (p<0.05).

**Supplemental Figure 3.**
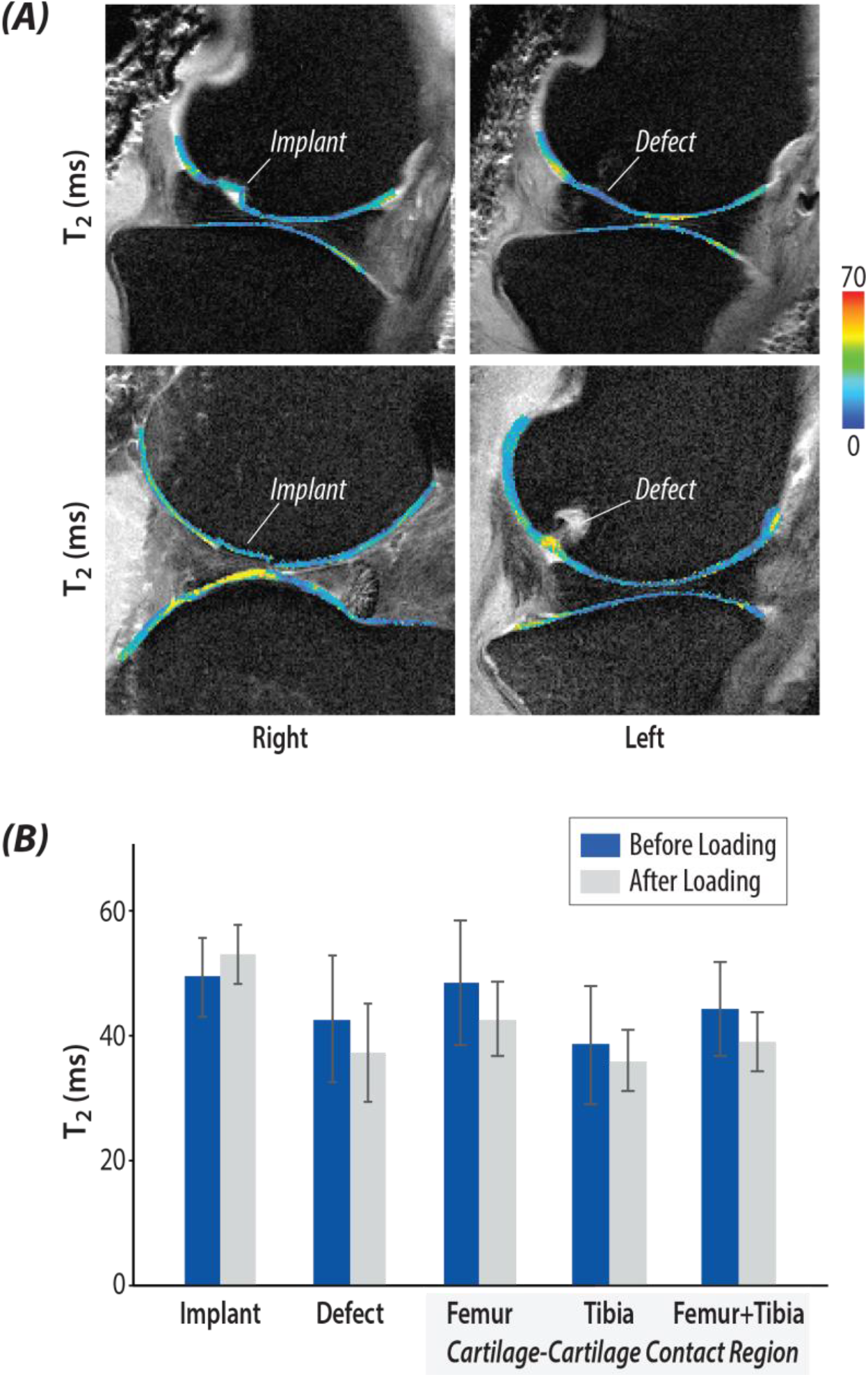
Relaxivity measurements of cartilage. (A) T_2_ mapping of implanted right knee and defect left knee used for evaluation of region-based differences. (B) No significant difference was found on average T_2_ value before and after loading in different ROI.

**Supplemental Figure 4.**
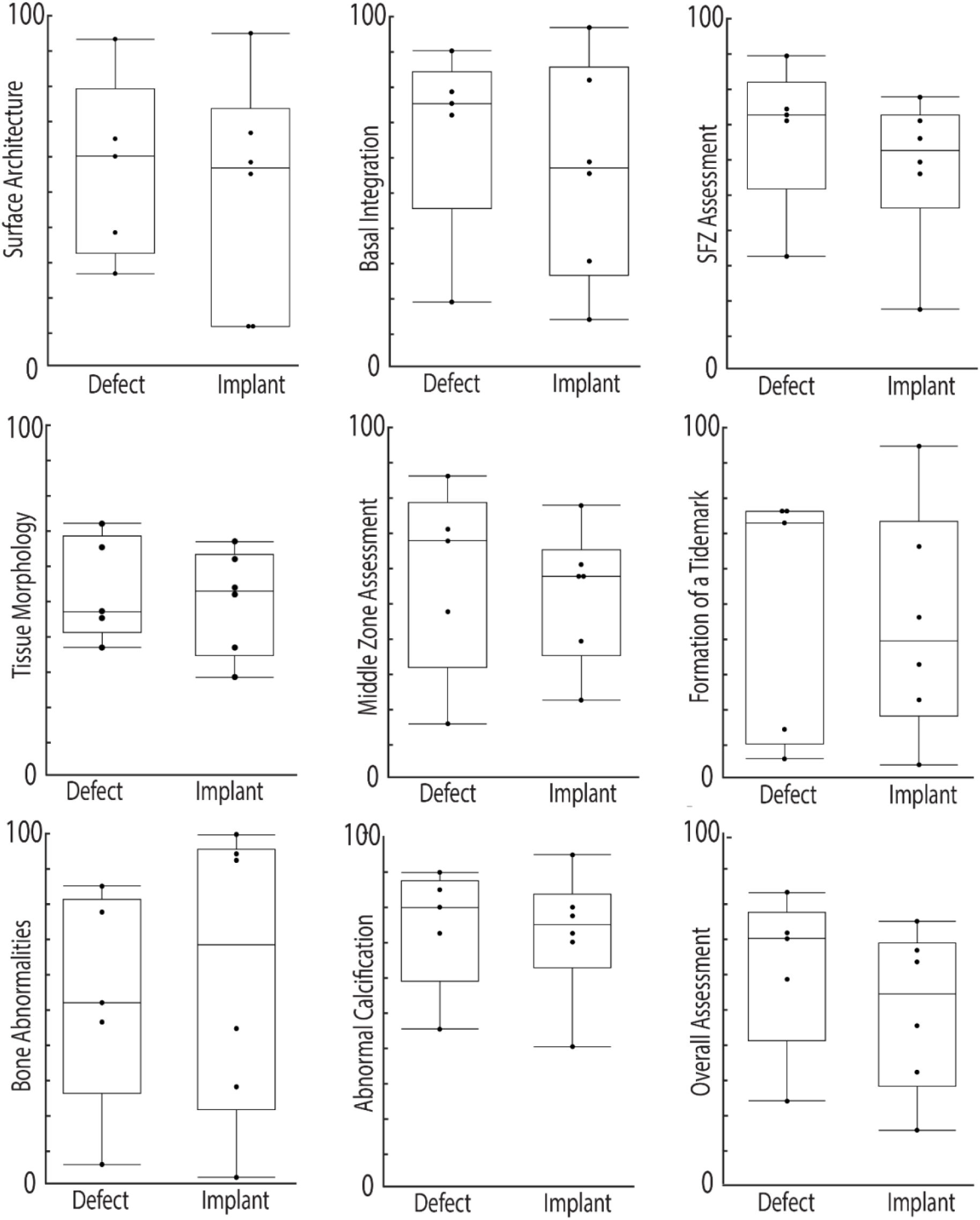
ICRS II Scoring. ICRS II visual histological score comparisons between the defect and implant groups. Scoring was completed by 3 blinded and independent observers. For detailed instruction on how images were scored, reference ICRS II guidelines ^52^. For all categories, a score of 100 indicates identical to native articular cartilage, where a score of 0 indicates no similarities to native cartilage.

**Supplemental Figure 5.**
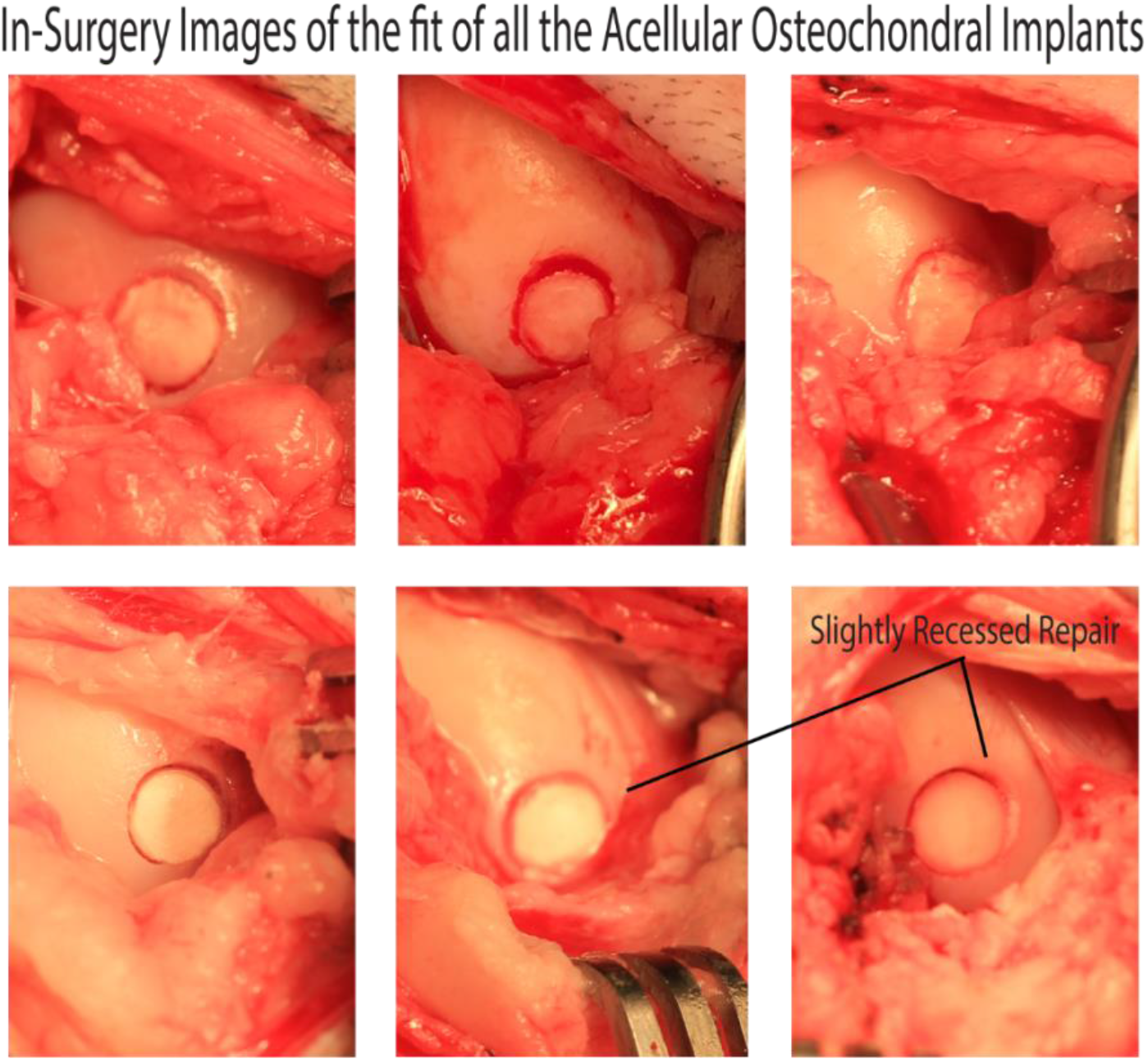
Surgical Fit of Acellular Implants. Surgical images of the acellular implant after the surgeon press-fit the construct into the defect. The surgeon placed the implant to try and align the cartilage surface, when possible.

